# A Phase Separation-Fortified Bi-Specific Adaptor for Conditional Tumor Killing

**DOI:** 10.1101/2023.07.19.549637

**Authors:** Yuyan Liu, Yuting Zhu, Weifan Xu, Pilong Li

## Abstract

A common approach in the development of therapeutic proteins is the use of synthetic ligands with multivalency, allowing for sophisticated control of signal transduction. Leveraging the emerging concept of liquid-liquid phase separation (LLPS) and its ability to organize cell surface receptors into functional compartments, we herein have designed modular ligands with phase-separation modalities to engineer programmable interreceptor communications and precise control of signal pathways, thus inducing the rapid, potent, and specific apoptosis of tumor cells. Despite their simplicity, these “triggers”, named phase-separated Tumor Killers (hereafter referred to as psTK), are sufficient to yield interreceptor clustering of death receptors (represented by DR5) and tumor-associated receptors, with the following features: LLPS-mediated robust high-order organization, well-choreographed conditional activation, and broad-spectrum capacity for potently inducing apoptosis of tumor cells. The development of novel therapeutic proteins with phase-separation modalities showcases the power of spatially reorganizing signal transduction. This approach enables the branching of cell fate and holds promising potential for targeted therapies against challenging tumors.

**One-Sentence Summary:** Engineered modular ligands with phase-separation modalities can drive lethal dialog with tumor cells.

## INTRODUCTION

The interplay between ligands and receptors has a profound influence on signal pathway transduction, shaping the efficiency, specificity, and outcome of cellular responses (Cuatrecasas, 1974). These interactions go beyond simple one-to-one relationships: they encompass diverse binding modes and receptor clustering, and thus determine cellular behavior and play a crucial role in cellular signaling networks (Angers *et al*, 2002; George *et al*, 2002; Manz & Groves, 2010; Reich *et al*, 1997; Schlessinger, 1988; Smith & Scott, 2002). Understanding the dynamic interplay between ligands and receptors is pivotal for unraveling the complexities of signal transduction and holds great potential for therapeutic interventions targeting signaling pathways.

Liquid-liquid phase separation (LLPS) has emerged as a key factor in the transduction of signals from cellular membrane receptor-based signaling pathways (Case *et al*, 2019; Ladbury *et al*, 2023; Shelby *et al*, 2023; Su *et al*, 2016). Through phase separation, biomolecules form distinct liquid-like clusters that create specialized microenvironments, enabling efficient signal transduction. These phase-separated assemblies regulate the spatial organization, concentration, and dynamics of membrane receptors, leading to enhanced receptor clustering, signaling activation, and downstream cellular responses (Chen *et al*, 2020; Shelby *et al*, 2021; Xiao *et al*, 2022). The influence of biomolecular phase separation on receptor signaling provides valuable insights into the regulation and coordination of complex cellular processes, thus paving the way for novel strategies to manipulate signaling pathways for therapeutic purposes.

Dysregulated signaling pathways resulting from specific ligand-receptor interactions contribute to various diseases, including cancer, autoimmune disorders, and regenerative disorders (Alexiou *et al*, 2010; Kim & Cochran, 2017). By exploiting the specificity of protein-protein interactions, protein-based therapeutics targeting specific ligand-receptor interactions show promise in combating these diseases (Leader *et al*, 2008). Protein engineering approaches, driven by rational and combinatorial design, enable the creation of proteins with desired properties. Furthermore, the advantages of ligand oligomerization in multi-receptor signaling complexes have gained attention (Courtney *et al*, 2009; Spencer *et al*, 1993). However, the controlled design of multivalent ligands with specific distances and topologies remains a challenge. Incorporating phase-separation properties in protein engineering strategies holds potential for modulating biological responses by altering natural ligand-receptor interactions.

As part of our research, we focused on developing a synthetic receptor system based on death receptor 5 (DR5, TNFRSF10B). DR5, a member of the tumor necrosis factor (TNF) receptor superfamily, triggers the extrinsic apoptotic pathway upon receptor multimerization (Holland, 2014). While DR5 has shown promise as a cancer therapeutic target, limitations such as drug-induced hepatotoxicity and inadequate receptor clustering have hindered clinical efficacy (Ichikawa *et al*, 2001). In our study, we utilized two ligand-receptor pairs: an anti-DR5 nanobody ligand with the DR5 receptor and a specific nanobody ligand with a tumor-specific antigen. These pairs act as orthogonal receptor-receptor communication channels, connected by phase separation elements, resulting in rapid and potent induction of tumor cell apoptosis with promising therapeutic potential.

## RESULTS

### Generation of psTK: A Platform for Phase Separation-Mediated Tumor Killing

Potency and selectivity are pivotal factors in the development of DR5 agonists. However, first-generation TNF-related apoptosis-inducing ligand (TRAIL) receptor agonists and mono-antibodies face limitations such as poor stability and impaired efficacy due to DR5 antagonist receptors. To overcome the limited valency of mono-antibodies, researchers introduced multivalent tandem DR5 nanobodies, which showed promise but raised concerns regarding hepatotoxicity **(Fig 1A)**. In the second generation, bi-specific antibodies targeting both DR5 and tumor-specific markers were introduced, yet the efficient activation of DR5 using high valency remained unexplored **(Fig 1A)**.

**Figure 1.**
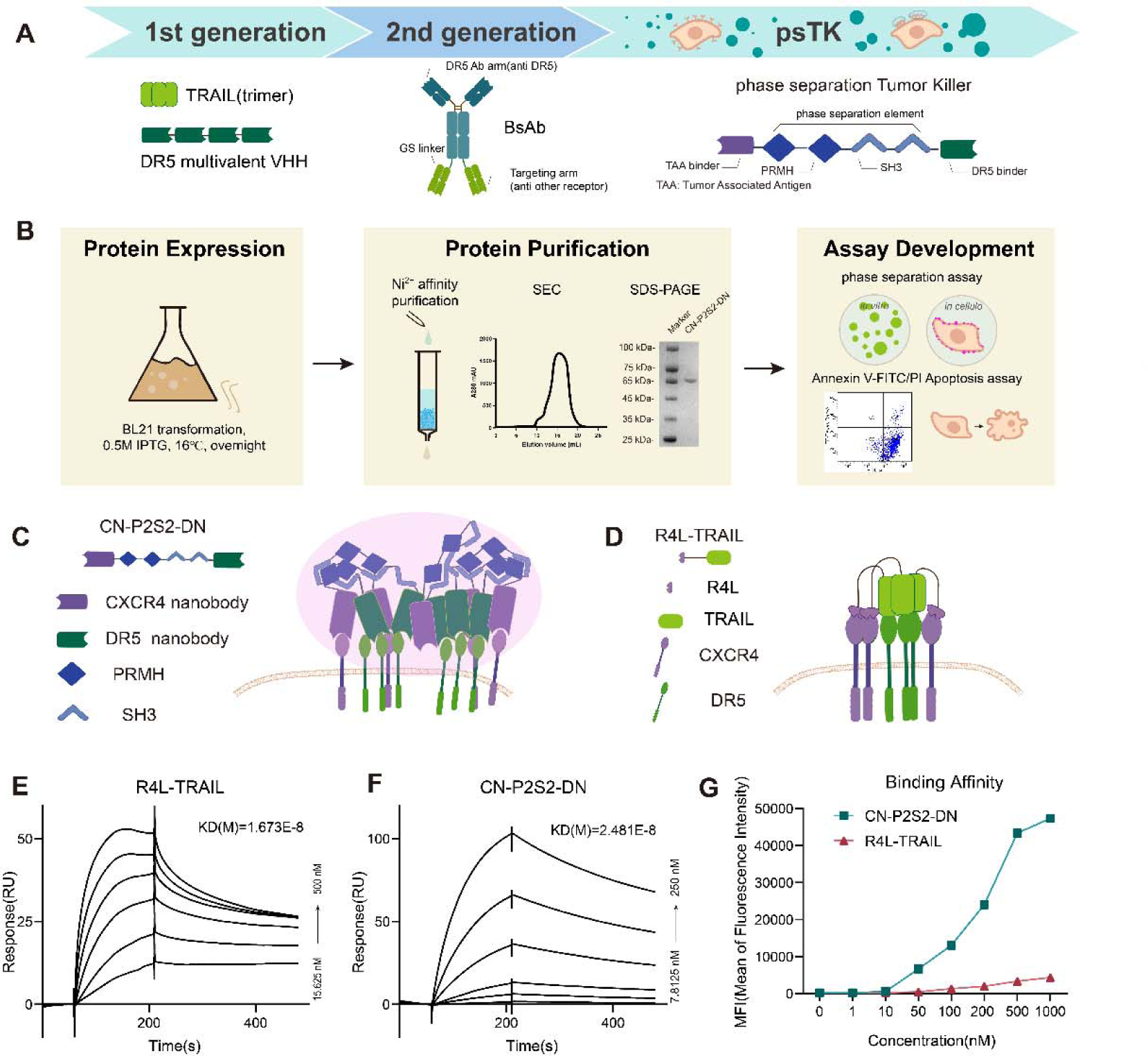
Platform for the Generation of Phase-Separating Tumor Killers (psTK). **A** Advancement of death receptor agonists as anti-cancer therapeutics. **B** Experimental workflow of the study. **C, D** Cartoon models of R4L-TRAIL and CN-P2S2-DN. **E, F** Surface plasmon resonance (SPR) affinity measurements. Sensorgrams depict the binding of R4L-TRAIL or CN-P2S2-DN to immobilized DR5 extracellular domain. **G** Binding affinity of R4L-TRAIL or CN-P2S2-DN to SJSA-1 tumor cells assessed by flow cytometry.

Driven by our commitment to enhance cancer therapeutics, we have developed the phase-separated Tumor Killer (psTK) platform **(Figs 1A and 1B)**. Our innovative approach involves engineering two receptors, CXC motif chemokine receptor 4 (CXCR4) and death receptor 5 (DR5), using a multivalent scaffold system with tandem repeats of (PRM)_2_ and (SH3)_2_ (Li *et al*, 2012), referred to as P2S2. This scaffold is combined with an agonist DR5 nanobody (DN) (Huet *et al*, 2014) and an antagonist CXCR4 nanobody (CN) (Jähnichen *et al*, 2010) to create a bi-specific ligand **(Fig 1C)**. Through gene synthesis, protein expression using a prokaryotic expression system, and purification, we achieved rapid and efficient production of these fusion proteins **(Fig 1B)**.

To compare our design with the second-generation approach, we also created R4L-TRAIL **(Figs 1D and EV1A)**, which incorporates TRAIL fused with a CXCR4-binding peptide(Liu *et al*, 2014). Surface plasmon resonance (SPR) results confirm the strong affinity of both R4L-TRAIL and CN-P2S2-DN proteins for DR5 **(Figs 1E and 1F, and EV1B-1C)**. Notably, CN-P2S2-DN exhibits markedly enhanced specific binding affinity in cellular assays compared to R4L-TRAIL and control proteins **(Figs 1G and EV1D)**.

By leveraging these designs, our aim is to optimize the potency and selectivity of DR5 activation, thereby paving the way for more effective and targeted cancer treatments.

### Cellular Effects of psTK: Phase Separation and Receptor Reorganization on the Cell Membrane

To gain insights into the cellular effects of psTK protein **(Fig EV2A)**, we conducted in vitro phase separation assays to confirm the LLPS properties. Confocal fluorescence microscopy was employed to visualize the behavior of the CN-P2S2-DN protein, aided by the molecular crowder PEG-8000. The results revealed that CN-P2S2-DN formed numerous spherical droplets, which settled onto the coverslip surface due to gravity **(Fig EV2B)**. Through time-lapse observations, we observed that these condensates underwent fusion and coalescence upon contact, forming larger droplets **(Fig EV2C; Movie EV1)**, confirming their liquid-like characteristics.

To further verify the liquid-like behavior, we performed fluorescence recovery experiments within a bleached region **(Fig EV2D; Movie EV1)**. The recovery of fluorescence demonstrated the free diffusion of CN-P2S2-DN protein molecules within the condensed phase, consistent with their liquid-like nature. These findings validate the LLPS properties of the CN-P2S2-DN protein and confirm its ability to undergo phase separation in vitro. The in vitro phase separation data provide a concentration threshold reference for subsequent cellular experiments. Based on these findings, we can proceed with the goal of conditionally activating psTK while ensuring the exclusive presence of CN-P2S2-DN without undergoing phase separation in solution.

Upon treating dual-positive DR5^+^/CXCR4^+^ SJSA-1 cells (Li *et al*, 2022), derived from osteosarcoma, with CN-P2S2-DN protein labeled with Alexa 546, we observed the self-association and aggregation of CN-P2S2-DN on the cell surface. These aggregates formed micron-sized puncta that had a propensity to fuse upon contact **(Figs 2A-2C)**. To gain a deeper understanding of the dynamic interactions between CN-P2S2-DN protein and receptors, we co-expressed CXCR4-mCherry and DR5-GFP in 293T cells and conducted real-time live-cell imaging after treating the cells with 200 nM CN-P2S2-DN. As expected, we observed the enrichment and colocalization of GFP and mCherry signals within the condensates formed by Alexa 647-labeled CN-P2S2-DN protein **(Figs 2D-2E)**. This indicates the ability of CN-P2S2-DN protein to cluster and induce intricate rearrangement of receptors. Importantly, in the absence of full-length CN-P2S2-DN, no colocalization or enrichment was observed (**Figs EV2E-2F**). Immunofluorescence analysis of endogenous receptors using recombinant proteins revealed a similar pattern of colocalization in SJSA-1 cells **(Figs 2F-2G, and EV2G-2H)**. It is noteworthy that the formation of puncta on the cell membrane required a concentration of only 200 nM CN-P2S2-DN, while in vitro LLPS required a significantly higher (micromolar) concentration of CN-P2S2-DN and addition of the crowder PEG-8000. We propose that the cell membrane receptors play a crucial role in this process, facilitating the binding and enrichment of CN-P2S2-DN. Once bound to the cell receptors, the phase separation backbone P2S2 has the opportunity to form condensates at a relatively low dose.

**Figure 2.**
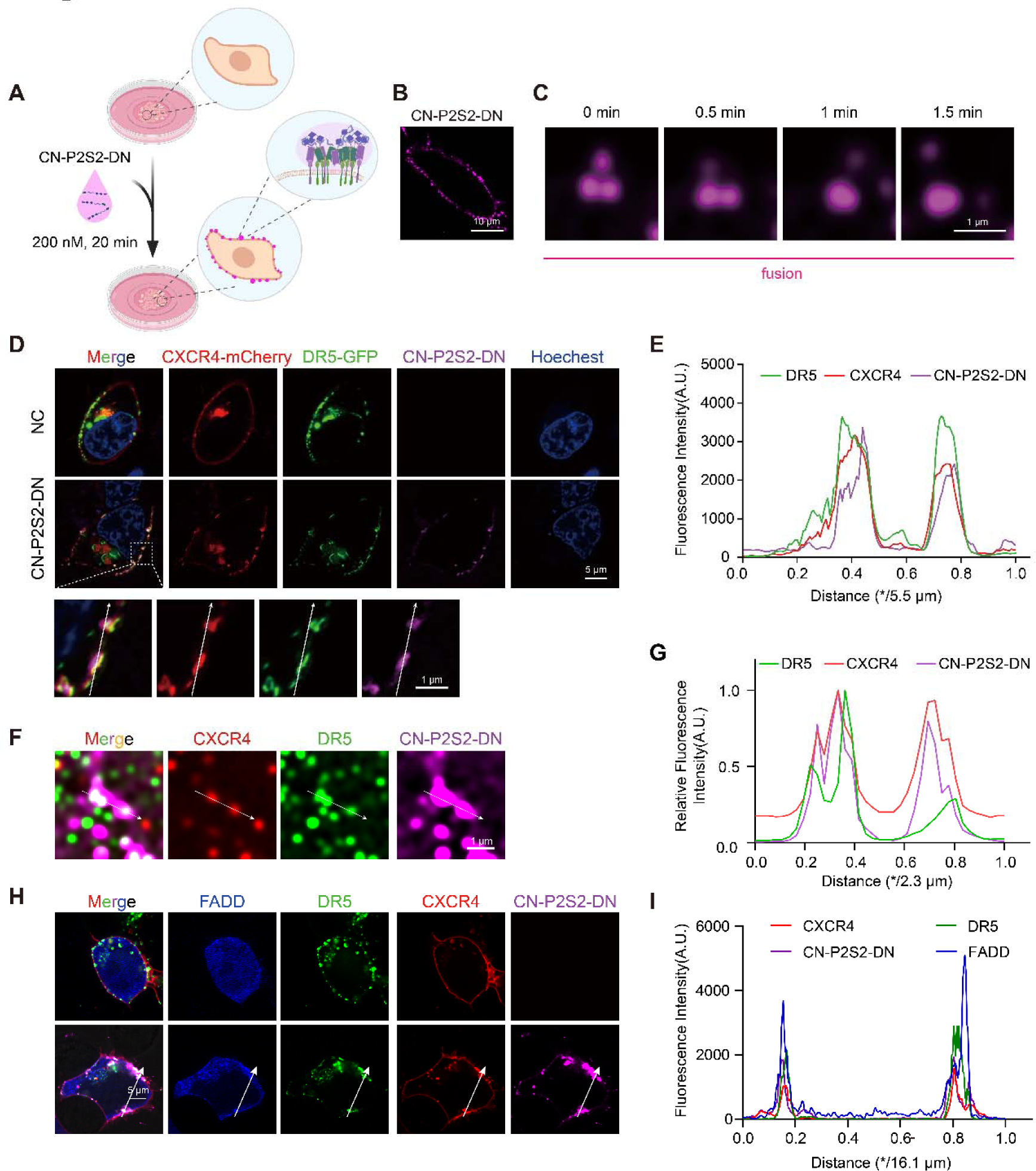
psTK induces phase separation and receptor reorganization on the cell membrane. **A** Diagram illustrating the method for incubating cells with the modular ligand CN-P2S2-DN. The ligand was labeled with Alexa 546. **B** Confocal fluorescence imaging of SJSA-1 cells treated with CN-P2S2-DN (200 nM). CN-P2S2-DN condensates formed on the cell membrane after 20 minutes of incubation. Scale bar, 10 μm. **C** Fusion of CN-P2S2-DN condensates on the cell. **D** HEK293T cells expressing DR5-EGFP and CXCR4-mCherry were treated with 200 nM CN-P2S2-DN (Alexa 647). Confocal fluorescence images are shown. **E** Colocalization analysis plot. Fluorescence intensities of CN-P2S2-DN, DR5- EGFP, and CXCR4-mCherry were quantified along the white arrows in the bottom panels of **(D)**. **F** Confocal imaging of immunofluorescence. SJSA-1 cells treated with 100 nM CN-P2S2-DN (Alexa 647) were fixed and incubated with the indicated primary and secondary antibodies. **G** Colocalization analysis plot. Fluorescence intensities of CN-P2S2-DN, DR5, and CXCR4 were quantified along the white arrows in **(F)**. **H** HEK293T cells expressing DR5-EGFP, CXCR4-mCherry, and FADD-BFP were treated with 200 nM CN-P2S2-DN (Alexa 647). Confocal fluorescence images are shown. **I** Colocalization analysis plot. Fluorescence intensities of CN-P2S2-DN, DR5- EGFP, CXCR4-mCherry, and FADD-BFP were quantified along the white arrows in **(H)**.

Further investigation into the dynamics of this system revealed fusion and fission events on the cell membrane through time-lapse observation (**Figs EV2I-2K**). SIM/STORM-TIRF observation revealed the time-lapse co-clustering of CN-P2S2-DN and DR5, providing further evidence of their interaction and condensate formation **(Fig EV2I)**. Moreover, Spatiotemporal analysis demonstrated the rapid redistribution of CN-P2S2-DN and CXCR4-mCherry from the unbleached area to the bleached area, while DR5-GFP exhibited a relatively solid-like property once recruited into the condensates containing these three components **(Figs EV2K; Movie EV2)**. Additionally, the overexpression of Fas-associating protein with a novel death domain (FADD), a downstream effector in the DR5 cascade signaling pathway (Sayers, 2011), resulted in its enrichment in the vicinity of the plasma membrane along with the DR5 condensates upon treatment with CN-P2S2-DN **(Figs 2H-2I)**. These findings highlight the dynamic behavior of the CN-P2S2-DN system and the colocalization of key components involved in the DR5 signaling pathway.

Based on these comprehensive results, we conclude that CN-P2S2-DN has the ability to drive the reorganization of cell membrane receptors through LLPS. These in vitro phase separation data provide valuable insights for subsequent cellular experiments, ensuring that sole CN-P2S2-DN is present without undergoing phase separation, thus achieving conditional activation upon contacting with the respective receptors.

### Selective Tumor Cell Death: Apoptosis Induction by psTK

We conducted cytotoxicity assays to investigate the potential of programming DR5- clustering-mediated apoptosis in specific cancer cell lines. SJSA-1 cells were exposed to CN-P2S2-DN and monitored in real time using a confocal microscope after staining with the apoptosis tracking dye YO-PRO. Compared to the control group, treatment with CN-P2S2-DN resulted in a series of significant morphological changes, including cell shrinkage and cellular content leakage **(Fig EV3A)**. Time-lapse images were captured **(Movie EV3)**, providing a dynamic visualization of the process. Cell death was observed as early as 90 minutes after the administration of 10 nM CN-P2S2-DN and reached a plateau after 3 hours in a time-and concentration-dependent manner **(Figs 3A-3B)**. To further evaluate apoptosis, FACS-based Annexin V/PI staining was performed. CN-P2S2- DN exhibited potent apoptosis-inducing activity in CXCR4^+^/DR5^+^ SJSA-1 and COLO 205 cells, with IC_50_ values of 10.1 nM and 28.9 nM, respectively **(Figs 3C and EV3B)**. As an example, in SJSA-1 cells, after a 3-hour treatment with CN-P2S2-DN at a concentration of 100 nM, 82.5% of the cells tested positive for Annexin V, whereas only 9.4% of CN-P2S2-DN-treated cells from the non-cancer line 293T showed Annexin V positivity, similar to the blank controls (4.6%) **(Figs 3C and EV3B)**. Intriguingly, CN-P2S2-DN exhibited comparable potency to the first-generation design, TRAIL **(Fig EV3C)**, and the second-generation design, R4L-TRAIL **(Fig EV3D)**.

**Figure 3.**
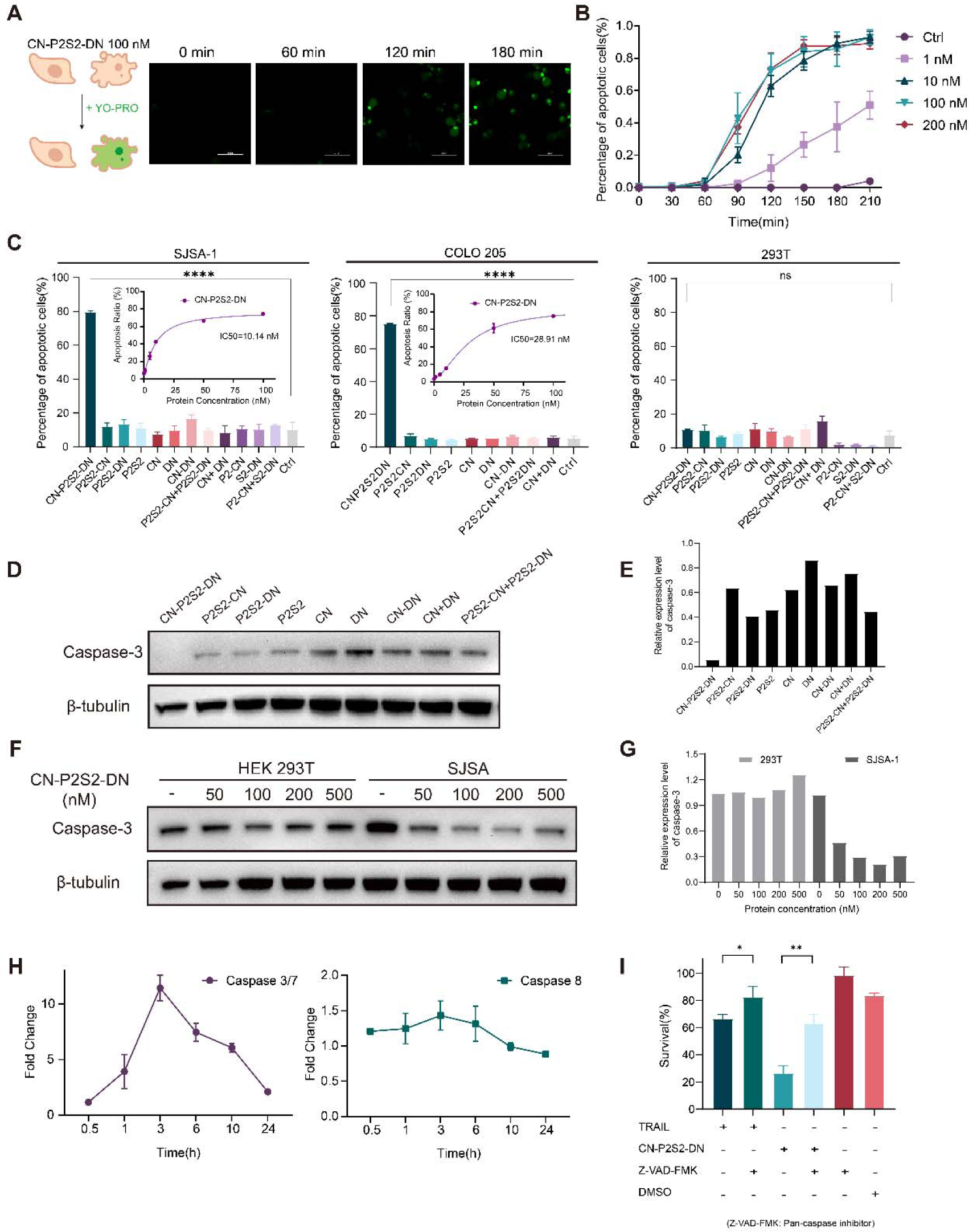
Selective Induction of Apoptosis in Tumor Cells by psTK. **A** Confocal live-cell imaging of SJSA-1 cells treated with 100 nM CN-P2S2-DN for 180 minutes. The cells were pre-incubated with the green fluorescent carbocyanine nucleic acid stain YO-PRO, which enters apoptotic cells through plasma membranes with increased permeability and binds to DNA. **B** Analysis of apoptosis ratio based on the images in **(A)**. **C** Annexin V/PI analysis of SJSA-1, COLO 205, and HEK293T cells treated with 100 nM CN-P2S2-DN or control proteins for 5 hours. Dose-response curves for CN-P2S2-DN are shown for SJSA-1 and COLO 205. Data are representative of three independent experiments. **D** Western blot analysis of caspase 3 activation in SJSA-1 cells upon treatment with 100 nM CN-P2S2-DN and control proteins. **E** Quantitative analysis of caspase-3/β-tubulin band intensity in **(D)**. **F** Western blot analysis of caspase 3 activation in HEK293T or SJSA-1 cells treated with different concentrations of CN-P2S2-DN. **G** Quantitative analysis of caspase-3/β-tubulin band intensity in **(F)**. **H** Kinetics of caspase 3/7 and caspase 8 activity, measured by Caspase-Glo® assay, following treatment of SJSA-1 cells with 100 nM CN-P2S2-DN. **I** Inhibition of CN-P2S2-DN-induced apoptosis in SJSA-1 cells by the pan-caspase inhibitor Z-VAD-FMK. TRAIL was used at a concentration of 0.1 nM and CN-P2S2-DN at 10 nM.

Western blot analysis revealed that CN-P2S2-DN treatment induced a significant specific and dose-dependent decrease in the level of full-length caspase 3, which is known to undergo cleavage-mediated activation during apoptosis **(Figs 3D-3G)**. Moreover, the activities of caspase 3/7 and caspase 8 in SJSA-1 cells increased over time, peaking at 3 hours of treatment with 100 nM CN-P2S2-DN **(Fig 3H)**. Further analysis using the pan-caspase inhibitor Z-VAD-FMK showed that DR5-mediated apoptosis was partially impaired, with a decrease of 16.1% by TRAIL and 36.8% by CN-P2S2-DN **(Fig 3I)**. This indicates a more sensitive response to CN-P2S2-DN treatment. Interestingly, expression of pAKT, an indicator of the CXCR4 signaling pathway associated with cell survival, displayed a “roller coaster” mode of expression fluctuations, suggesting the activation of CXCR4 as a last-ditch struggle of tumor cells **(Figs EV3E-3F)**. These findings collectively show that CN-P2S2-DN effectively induces apoptosis in specific cancer cell lines through DR5 clustering, and demonstrate its potential as a therapeutic agent.

### Harnessing Selectivity and Dynamics: Valency-and Time-Dependent Synthetic Apoptosis with psTK

After demonstrating that CN-P2S2-DN triggers conditional apoptosis in cellular models, we aimed to elucidate the underlying mechanism of action. Flow cytometry measurements provided further confirmation of higher expression levels of CXCR4 and DR5 in tumor cells compared to 293T cells **(Fig 4A)**. To investigate the impact of DR5 expression levels on CN-P2S2-DN-induced cell apoptosis, we conducted experiments involving DR5 knockdown and overexpression. The results showed a significant inhibition of apoptosis rate upon DR5 knockdown, while overexpressing DR5 did not yield substantial differences in the potency of CN-P2S2-DN **(Figs 4B-4C)**. These findings suggest that CN-P2S2-DN may efficiently aggregate DR5 into phase-separated condensates even in its natural state, leading to the saturation of its signaling pathway.

**Figure 4.**
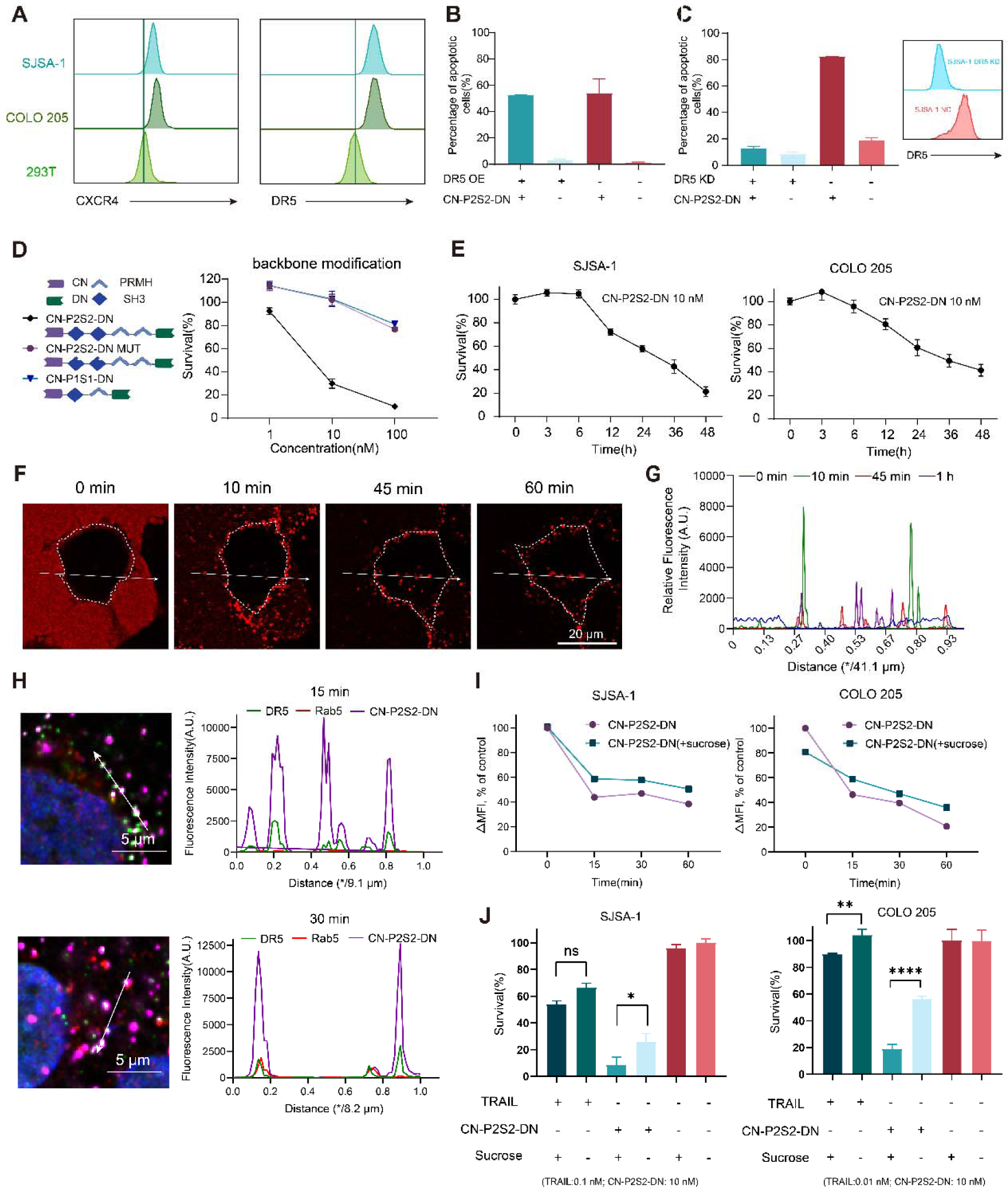
Selective, Valency-Dependent, and Time-Dependent Induction of Apoptosis by psTK. **A** Flow cytometry analysis of surface CXCR4 and DR5 receptor expression in SJSA-1, COLO 205, and HEK293T cells, highlighting the differential receptor levels among the cell lines. **B** Overexpression of DR5 does not alter the sensitivity of SJSA-1 cells to 100 nM CN-P2S2-DN treatment. **C** Knockdown of DR5 reduces the sensitivity of SJSA-1 cells to 100 nM CN-P2S2- DN treatment. **D** Backbone modifications affect the cytotoxicity of psTK. Disruption of the PRM-SH3 interaction, reduction of valency, or changes in the module positions attenuate the anti-tumor activity of psTK in SJSA-1 cells. **E** Time-dependent tumor-killing function of CN-P2S2-DN. Survival analysis of SJSA-1 and COLO 205 cells exposed to 10 nM CN-P2S2-DN for varying durations. **F** Time-lapse confocal fluorescence imaging of SJSA-1 cells treated with 100 nM CN-P2S2-DN over a 60-minute period. From 45 minutes onwards, CN-P2S2-DN is observed entering the cytosol (white arrows). **G** Measurement of relative fluorescence intensity over time. The plot shows the dynamic distribution and intensity of the fluorescence signal of the recombinant protein CN-P2S2-DN in cells at different time points, as indicated by white arrows in **(F)**. **H** Representative immunofluorescence imaging of endogenous DR5 and Rab5 localization in SJSA-1 cells after treatment with 100 nM CN-P2S2-DN for 15 minutes or 30 minutes. At 30 minutes, colocalization of DR5 and CN-P2S2-DN with Rab5 is observed (indicated by the white arrow). **I** Time-dependent decrease in surface expression of DR5 by CN-P2S2-DN, as determined by flow cytometry. Hypertonic sucrose inhibits CN-P2S2-DN-induced endocytosis of DR5 in SJSA-1 and COLO 205 cells. Cells were pre-incubated with or without 250 mM sucrose for 1 hour prior to treatment with 10 nM CN-P2S2-DN. **J** Hypertonic sucrose pretreatment sensitizes SJSA-1 and COLO 205 cells to treatment with CN-P2S2-DN (10 nM) or TRAIL (0.1 nM for SJSA-1, 0.01 nM for COLO 205).

We also explored the role of multivalent interactions in mediating phase separation. By reducing the valency of the phase separation scaffold protein and introducing mutations (specifically proline to alanine (Kurochkina & Guha, 2013)) to disrupt the interplay between multivalent domains, we highlighted the critical role of multivalency-mediated phase separation as a key factor driving conditional apoptosis in tumor cells **(Fig 4D)**. Notably, the well-designed quadrivalent scaffold of P2S2 demonstrated remarkable efficacy in triggering apoptosis, compared to the poor potency observed in the dose-response measurement of CN-P1S1-DN **(Fig EV4A-4D)**.

Shifting our focus to the temporal aspect, we were intrigued to discover that even when cells were treated with a lower concentration of CN-P2S2-DN (10 nM), they still exhibited a time-dependent and persistent death response, as evidenced by CellTiter-Glo (CTG) assays at the indicated time points. The onset of this time-dependent cell death was observed around 6 hours after treatment, indicating the progressive nature of the apoptotic process **(Fig 4E)**. Further detailed observations using confocal microscopy revealed that as early as 10 minutes after adding CN-P2S2-DN to the cell culture medium, significant focal accumulations were observed at the cell membrane **(Figs 4F- 4G; Movie EV4)**. Efficient ligand binding facilitated the rapid internalization of DR5, as indicated by its colocalization with Rab5, an early endosome marker **(Fig 4H and EV4E)**. Flow cytometry analysis confirmed the occurrence of receptor internalization, with the cell surface expression of DR5 was significantly reduced to 38.3% and 20.6% in SJSA-1 and COLO 205 cells, respectively, after 60 minutes of CN-P2S2-DN treatment **(Fig 4I)**. Interestingly, pretreatment with the pan-endocytosis inhibitor sucrose (Guo *et al*, 2015) effectively hindered the internalization process, resulting in an upregulation of surface DR5 expression to 50.5% and 35.8% in SJSA-1 and COLO 205 cells, respectively **(Fig 4I)**. Moreover, inhibiting DR5 endocytosis through sucrose treatment significantly enhanced the apoptotic induction capability **(Figs 4J and EV4F-4G)**. These findings suggest that inhibiting DR5 internalization can effectively increase the sensitivity of cells to the protein and enhance their apoptotic response. In summary, the multivalent interactions, time-dependent response, and inhibition of DR5 endocytosis contribute to the efficient induction of conditional apoptosis by CN-P2S2-DN in tumor cells.

### Expanding the Reach of psTK: Targeting Additional Tumor Cells and Diverse Tumor Targets

To verify the compatibility and effectiveness of our system across multiple cancer indications **(Fig 5A)**, we evaluated CN-P2S2-DN-mediated cytotoxicity in a panel of cell lines from different cancer types **(Fig 5B)**. Relative expression levels of CXCR4 and DR5 were determined, and distinct responses to multivalent ligand treatment were observed among the different cell lines **(Fig 5C and EV5A-5C)**. High-responder cell lines, including colorectal, sarcoma, lung, and prostate cancer cell lines, exhibited a more potent lethal ratio (Ratio_min_ > 40%) **(Fig 5D)**. More data on orthogonal combinations of synthetic ligands with tumor cell lines are provided in **Fig EV5D**. The NCI-H1975 cell line showed the maximum response with a survival ratio of 2.8% **(Fig 5D)**. In the case of NCI-H1975, the potent induction of cytotoxicity was surprising, given that these cells have a DR5:CXCR4 ratio much higher than 1 with regard to the in-silico analyses from DepMap database. This finding suggests a potential advantage in overcoming tumor escape mechanisms associated with low target antigen expression.

**Figure 5.**
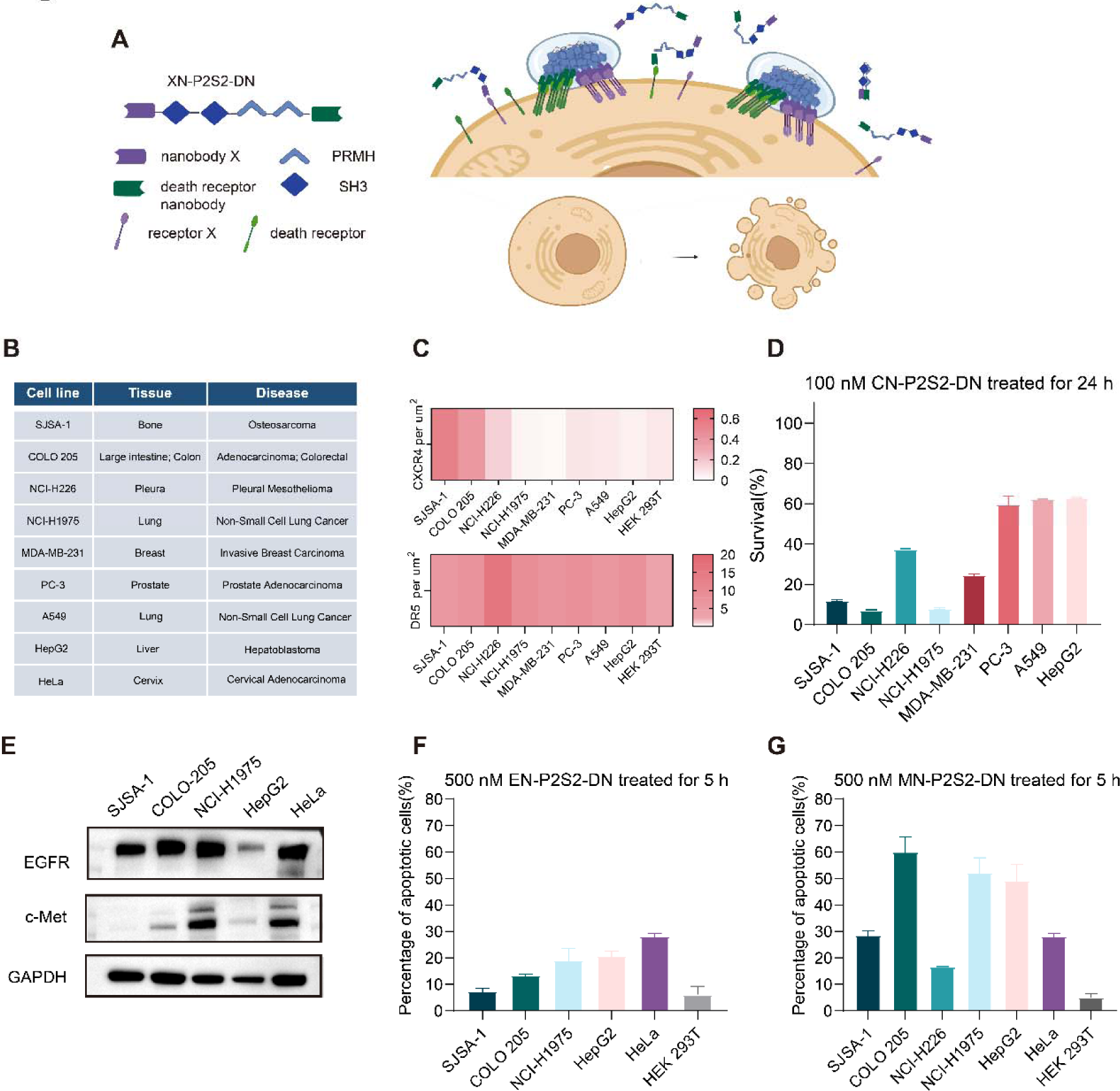
Expansion of the psTK Platform: Evaluation of Distinct Targets in Tumor Cell Lines. **A** Diagram illustrating the expansion of the psTK platform. **B** Information regarding several tumor cell lines tested in this study. **C** Flow cytometry analysis of relative surface expression of CXCR4 and DR5 in the tested cell lines. **D** Cell survival analysis of various tumor cell lines after 24 hours of treatment with 100 nM CN-P2S2-DN. Survival was assessed using the CellTiter-Glo® assay. **E** Western blot analysis of EGFR and c-Met expression in the tested cell lines. **F** Cell survival analysis of various tumor cell lines after 5 hours of treatment with 500 nM EN-P2S2-DN. Apoptosis was assessed using the Annexin V-FITC/PI staining. **G** Cell survival analysis of various tumor cell lines, assessed using the Annexin V-FITC/PI staining, after 5 hours of treatment with 500 nM MN-P2S2-DN.

Furthermore, we designed other ligands based on a similar framework **(Fig 5A)** to induce alternative types of interreceptor clustering with the same apoptotic output. c-Met (also known as hepatocyte growth factor receptor, HGFR) and epidermal growth factor receptor (EGFR) are highly expressed tumor-associated antigens (TAAs). As described above, we engineered synthetic ligands for these promising cancer targets: EN-P2S2-DN for EGFR and MN-P2S2-DN for c-Met **(Fig EV5A)**. After verifying the expression patterns of the respective targets via *in silico* prediction and western blot analysis **(Figs 5E**, and **EV5E-5F)**, we performed cytotoxicity assays. Treatment with EN-P2S2-DN against EGFR, a target extensively evaluated in the clinic, also exhibited good performance in NCI-H1975, an EGFR mutant-positive cell line **(Fig 5F)**. Intriguingly, c-Met showed a distinct advantage as a pan-cancer target, as reflected by the excellent apoptosis-inducing effect of the ligand MN-P2S2-DN on the majority of cell lines, similar to the effect of CN-P2S2-DN **(Fig 5G)**. In conclusion, our findings demonstrate the broad applicability of XN-P2S2-DN in inducing conditional apoptosis across multiple cancer types, with distinct responses observed among different cell lines. Furthermore, the design of alternative ligands targeting different TAAs highlights the potential for personalized and targeted cancer therapeutics.

## DISCUSSION

In summary, we have successfully developed synthetic structures using our minimal logic to control cell fate, specifically apoptosis. Our strategy utilizes phase separation-mediated inter-receptor clustering and communication to activate death receptor signaling. This innovative approach allows us to manipulate cellular processes and selectively induce programmed cell death. The remarkable diversity, flexibility, and robustness exhibited by these structures emphasize the programmable potential on signaling receptors. Across all of our systems, we have observed conditional activation events that lead to “programmed” cell death specifically in tumor cells. This activation occurs only when multivalent ligands bind to DR5 and a specific TAA, triggering the unfolding of “self-locking” structures through phase separation mechanisms. These findings have great significance, as they provide powerful tools for systematically manipulating multi-receptor complexes and modulating signaling outputs. This creates tremendous opportunities for targeted treatments of various diseases, and opens up a promising avenue for the field of precision medicine.

Previous efforts in engineering TRAIL death receptor agonists (TDRAs) have primarily concentrated on two approaches: recombinant TRAIL and multimeric anti-DR5 agonistic antibodies. Upon its discovery, TRAIL was initially believed to possess the ability to selectively induce apoptosis in tumor cells while sparing normal cells (Wiley *et al*, 1995). This promising property sparked a significant amount of scientific research aimed at exploring TRAIL-based anti-cancer strategies. However, during the early attempts to develop TDRAs for cancer therapy, numerous challenging technical problems arose. Recombinant forms of TRAIL, such as Dulanermin, have exhibited disappointing clinical results due to their short half-life and the intrinsic resistance of primary tumor cells (Von Karstedt *et al*, 2017). Furthermore, non-apoptotic signaling pathways activated by TRAIL decoy receptors (TRAIL-R3, TRAIL-R4, and OPG) have the inevitable effect of negatively regulating apoptosis. Although multivalent antibodies have shown greater specificity and higher valency compared to conventional antibodies (Huet *et al*., 2014; Wang *et al*, 2021), the phase Ⅰ clinical trial was suspended due to acute liver toxicity concerns (Camidge *et al*, 2010). Taking these limitations into account, the second generation of TDRA system design has made significant progress. One of the second-generation approaches involves the use of bivalent antibodies that can target two different epitopes, thereby improving specificity and reducing off-target toxicity. Roche has developed a bi-specific antibody (BsAb) called RG7386, which has been designed to simultaneously engage DR5 on tumor cells and fibroblast-activation protein (FAP) on tumor stroma fibroblasts (Brünker *et al*, 2016). Similarly, another study has designed the BACA-1 antibody to target FOLR1 and DR5 receptors for ovarian cancer (OvCa) therapy (Shivange *et al*, 2018). Overall, the challenge of balancing selectivity and efficacy remains an ongoing obstacle, and necessitates the emergence of more innovative molecular entities.

Herein, our innovative modular platform technology allows us to expand beyond the tedious and intricate design and manufacturing limitations to some extent. Compared to previously reported TDRAs, our phase separation-based platform offers significant advantages in terms of expedited expression, bioactivity testing, and matrix screening of bi-specific binders. Through the use of our psTK system, we have made several unexpected discoveries. First, psTK represents a departure from previous phase separation-based fusion protein designs (Li *et al*., 2022). The unique “self-locking” structural configuration of the modular ligand ensures that its phase separation-mediated enrichment occurs efficiently when it simultaneously binds to both types of receptors. This is evident from the weak in vitro phase separation ability of the ligand, which requires the presence of crowding reagents, as well as the ability of the ligand to co-cluster cell membrane surface receptors at extremely low concentrations. These data strongly indicate a unique advantage of the psTK design: there is a significant reduction in the effective threshold concentration of ligand required for activity, and a simultaneous increase in the stability of the phase-separated ligand. Furthermore, the corresponding forms of ligand combinations are also more diverse, and not limited to the nanobody used in this study. Other applicable options include scFv, artificially synthesized binders, and natural ligands. Considering the increasing number of studies, including this one, confirming that membrane receptor-based phase separation contributes to the regulation of downstream signaling pathways, suggesting that it might be sufficient for synthetic ligands to possess binding activity, without stringent criteria of activation or inhibition.

What is particularly intriguing is that, unlike the traditional “one-to-one” or “many-to-one” binding modes of therapeutic proteins, the receptor clustering phenomenon triggered by phase separation enables an effective “one-to-many” collective effect. This discovery opens up new possibilities for therapeutic interventions. Harnessing the power of receptor aggregation through phase separation allows for a more efficient and effective approach to target multiple receptors simultaneously. In addition, it offers a distinct advantage in addressing difficult targets with low expression levels or mutations that impact binding in the extracellular regions.

Not limited to apoptosis, this platform shall demonstrate its multidimensional applications in downstream signaling responses, encompassing proliferation, differentiation, cytokine secretion, cytotoxicity, and other events. Moreover, unlike dual-antibody development platforms that primarily focus on optimizing the production methods for bi-specific antibodies, this platform technology holds the potential for rapid, efficient, and precise discovery of novel combinations of targets by bi-specific ligands, effectively addressing the significant scarcity of new target combinations. This is achieved through a flexible and controllable multivalent phase separation system that anchors combinations of cell surface proteins, inducing membrane receptor phase separation, and further investigating the cellular phenotypic changes and biological functions corresponding to specific protein combinations. Currently, we have successfully screened and identified entirely new and specific combinations of receptors, which are aberrantly activated in tumors, and the co-clustering leads to either selective elimination of tumor cells or efficient degradation of membrane receptor proteins (data not shown). Overall, while the full potential of the phase-separated Engineering Therapeutic Protein (psETP) platform, represented by psTK, is still being established, we anticipate that this technology, when combined with modular protein engineering, will provide additional opportunities for manipulating membrane receptor function. This has significance for both fundamental research and the exploration of potential therapeutic applications.

## MATERIALS AND METHODS

### Materials

#### HEK293T cell line

Human kidney epithelial cell line HEK293T was maintained in DMEM (HyClone, Cytiva) plus 10% fetal bovine serum (GIBCO, Thermo Fisher Scientific), and 100 Units/ml Pen-Strep (GIBCO, Thermo Fisher Scientific), at 37°C, 5% CO_2_.

#### SJSA-1 cell line

Human Osteosarcoma cell line SJSA-1 (formerly OsA-CL) was maintained in RPMI-1640 (GIBCO, Thermo Fisher Scientific) plus 10% fetal bovine serum (GIBCO, Thermo Fisher Scientific), and 100 Units /ml Pen-Strep (GIBCO, Thermo Fisher Scientific), at 37°C, 5% CO_2_.

#### COLO 205 cell line

Human Colon Adenocarcinoma cell line COLO 205 was maintained in RPMI-1640 (GIBCO, Thermo Fisher Scientific) plus 10% fetal bovine serum (GIBCO, Thermo Fisher Scientific), and 100 Units /ml Pen-Strep (GIBCO, Thermo Fisher Scientific), at 37°C, 5% CO_2_.

#### NCI-H226 cell line

Human lung squamous cell carcinoma cell line NCI-H226 was maintained in RPMI-1640 (GIBCO, Thermo Fisher Scientific) plus 10% fetal bovine serum (GIBCO, Thermo Fisher Scientific), and 100 Units/ml Pen-Strep (GIBCO, Thermo Fisher Scientific), at 37°C, 5% CO_2_.

#### NCI-H1975 cell line

Human lung squamous cell carcinoma cell line NCI-H1975 was maintained in RPMI-1640 (GIBCO, Thermo Fisher Scientific) plus 10% fetal bovine serum (GIBCO, Thermo Fisher Scientific), and 100 Units/ml Pen-Strep (GIBCO, Thermo Fisher Scientific), at 37°C, 5% CO_2_.

#### MDA-MB-231 cell line

Human mammary gland epithelial cell adenocarcinoma cell line MDA-MB-231 was maintained in F-12K (GIBCO, Thermo Fisher Scientific) plus 10% fetal bovine serum (GIBCO, Thermo Fisher Scientific), and 100 Units/ml Pen-Strep (GIBCO, Thermo Fisher Scientific), at 37°C, 5% CO_2_.

#### PC-3 cell line

Human prostate adenocarcinoma cell line PC-3 was maintained in DMEM (HyClone, Cytiva) plus 10% fetal bovine serum (GIBCO, Thermo Fisher Scientific), and 100 Units/ml Pen-Strep (GIBCO, Thermo Fisher Scientific), at 37°C, 5% CO_2_.

#### A549 cell line

Human lung Carcinoma cell line A549 was maintained in DMEM (HyClone, Cytiva) plus 10% fetal bovine serum (GIBCO, Thermo Fisher Scientific), and 100 Units /ml Pen-Strep (GIBCO, Thermo Fisher Scientific), at 37°C, 5% CO_2_.

#### HepG2 cell line

Human liver hepatocellular carcinoma cell line HepG2 was maintained in MEM (GIBCO, Thermo Fisher Scientific) plus 10% fetal bovine serum (GIBCO, Thermo Fisher Scientific), and 100 Units/ml Pen-Strep (GIBCO, Thermo Fisher Scientific), at 37°C, 5% CO_2_.

#### HeLa cell line

Human cervical carcinoma cell line HeLa was maintained in DMEM (HyClone, Cytiva) plus 10% fetal bovine serum (GIBCO, Thermo Fisher Scientific), and 100 Units/ml Pen-Strep (GIBCO, Thermo Fisher Scientific), at 37°C, 5% CO_2_.

### Methods

#### Protein expression and purification

The recombinant proteins were overexpressed in *E. coli* BL21 (DE3). After overnight induction by 0.5 mM isopropyl β-D-thiogalactoside (IPTG) at 16°C in LB medium, cells were harvested and suspended in the buffer: 40mM Tris-HCl (pH 7.4), 500mM NaCl, and 10mM imidazole. After cell lysis and centrifugation, the recombined proteins were purified to homogeneity over the HisTrap column and eluted with a linear imidazole gradient from 20 mM to 500 mM. The proteins were further purified sequentially by MBPTrap HP columns and size-exclusion chromatography using a Superdex 200 Increase 10/300 GL column (Cytiva) in elution buffer (40 mM HEPES, pH7.5, 500mM NaCl, 5% glycerol).

#### Surface Plasmon Resonance (SPR)

Surface plasmon resonance measurements were conducted at Center of Pharmaceutical Technology, Tsinghua University, using the Biacore^TM^ S200 biosensor (Cytiva). Purified DR5-ECD was immobilized on a CM5 sensor chip and the binding experiments were run using single-cycle kinetics at 25 °C with the running buffer (40 mM HEPES pH 7.5, 500 mM NaCl). CN-P2S2-DN protein was injected for 150 s at a flow rate of 30 µL/min with a dissociation time of 270 s, using corresponding concentrations of CN-P2S2-DN (1/10 serial dilutions from 1 to 1,000 nM). The data were processed using Biacore S200 Evaluation Software Version 1.0, and fit to a Steady State Affinity model to calculate the equilibrium dissociation constant (KD).

#### Triton X-114-assisted Endotoxin removal

Endotoxin removal from protein solutions was performed as previously described with slight modifications. Briefly, Triton X-114 (Sigma-Aldrich) was added to the protein solution to a final Triton X-114 concentration of 1% v/v. The solution was incubated at 4°C for 60 min with constant stirring. Subsequently, the sample was transferred to a water bath set at 30°C and incubated for 30 min followed by centrifugation at 20,000 g for 20 min at room temperature. The upper part containing the protein was separated from the Triton X-114 layer and transferred into endo-free tubes by pipetting.

#### Protein labeling

Recombinant proteins were labeled by incubating with a 1:1 molar ratio of Alexa Fluor™ 647 C_2_ Maleimide or 488 C_5_ Maleimide or 546 C5 Maleimid (Thermo Fisher Scientific) for 1 h at room temperature with continuous stirring. The free dyes were removed by centrifugation in Zeba™ Spin Desalting Columns (Thermo Fisher Scientific, 89882), and the labeled proteins were stored at -80°C.

#### Phase separation assay

*In vitro* LLPS experiments were performed at room temperature. All samples were seeded and recorded on 384 low-binding multi-well 0.17 mm microscopy plates (In Vitro Scientific) and sealed with optically clear adhesive film. Recombinant proteins were diluted to the indicated final concentrations in the reaction buffer (20 mM HEPS, pH 7.4, 150 mM NaCl, and 10% PEG8000) with a total volume of 10 µl to induce phase separation.

For *in cellulo* assays, cells were seeded into 4-well chamber 35 mm dishes with a density of 5 × 10^5^ cells/well. With treatment of recombinant proteins at 37°C for indicated time, imaging was performed with a NIKON A1R HD25 confocal microscope equipped with a 100× oil immersion objective.

#### Immunofluorescence

Cells on coverslips were treated with the respective proteins at the designated concentration. At the indicated time points, the cells were fixed in 4% paraformaldehyde, permeabilized in 0.1% TritonX-100, blocked in 0.5% BSA/PBS, and incubated with primary antibodies overnight at 4°C, followed by secondary antibodies for 1 hr. The cells were then sealed in 4′,6′-diamidino-2-phenylindole (DAPI) (Beyotime), and examined with a NIKON A1R HD25 confocal microscope.

#### Flow Cytometry-Based Protein Binding Assay

Protein binding affinity was measured using flow cytometry. 200,000 cells were harvested and washed with PBS+1%FBS. Then the protein with corresponding concentration was incubated with the cells for 30 minutes in 37℃ at volume of 100 μl. After washing with PBS+1%FBS, incubating the cells with APC anti-His Tag Antibody (BioLegend) for 20 minutes on ice. Lastly, washing the cells with PBS+1%FBS and resuspend the pellet with 200 μl culture medium. The sample was then examined by BD LSRFortessa SORP flow cell sorter.

Protein binding affinity was assessed using a flow cytometry-based method. For each assay, 200,000 cells were harvested and washed with PBS containing 1% FBS. The cells were then incubated with the protein of interest at the appropriate concentration for 30 minutes at 37°C in a total volume of 100 μl. Following incubation, the cells were washed with PBS containing 1% FBS and incubated with APC anti-His Tag Antibody (BioLegend) for 20 minutes on ice. Subsequently, the cells were washed again with PBS containing 1% FBS and resuspended in 200 μl of culture medium. The samples were then analyzed using a BD LSRFortessa SORP flow cytometer.

#### Cell Surface Protein Expression Analysis

For the analysis of cell surface protein expression, 200,000 cells were harvested and washed with PBS+1%FBS. The cells were then incubated with the corresponding antibodies (typically at a dilution of 1:20) for 20 minutes on ice in 50 μl of culture medium. Subsequently, the cells were washed with PBS+1%FBS and resuspended in 200 μl of culture medium. Flow cytometric analysis was performed using a BD LSRFortessa SORP instrument. The precise surface receptor expression level, measured as the number of antibodies bound per cell, was calculated using a PE Fluorescence quantitation kit.

#### Cell Viability Assay

Cells were seeded in 96-well plates and allowed to incubate overnight for 16 hours at 37°C. Following the incubation, the cells were treated with the respective recombinant proteins for the indicated time points at 37°C. To assess cell viability, the CellTiter-Glo® Luminescent Cell Viability Assay (Promega) was employed, following the manufacturer’s instructions. Luminescence signals were measured using a SPARK luminometer (TECAN).

For the caspase inhibition assay, cells were pre-treated with 10 μM Z-VAD-FMK for 1 hour prior to the addition of CN-P2S2-DN.

To assess the role of endocytosis, cells were pre-incubated with 250 mM sucrose for 60 minutes and then transferred to regular culture medium before the treatment with CN-P2S2-DN.

These assays were conducted to investigate cell viability, the impact of caspase inhibition, and the involvement of endocytosis in response to CN-P2S2-DN treatment.

#### Flow cytometry-based apoptosis detection

Cells (1×10^6^) were treated with endotoxin-free proteins with the designated concentrations. After incubation for indicated time points at 37°C, flow cytometric analyses of FITC Annexin V staining (Beyotime) were proceeded to measure apoptosis, according to the manufacturer’s instructions.

#### Caspase Activity Assay

The Caspase-Glo assays were utilized to assess the activity of caspase-3/7 or caspase-8 in response to CN-P2S2-DN treatment. SJSA-1 cell (10,000 cells/well) were seeded in half-area 96-well plates and incubated overnight at 37°C. After the incubation period, the cells were treated with 100 nM CN-P2S2-DN for the indicated times. At the endpoint, the Caspase-Glo 3/7® or Caspase-Glo 8® assay (Promega) was performed according to the manufacturer’s protocol. Luminescence signals were measured using a SPARK luminometer (TECAN).

#### Live cell analysis

Cells were seeded into 4-well chamber 35 mm dishes at a density of 5 × 10^5^ cells/well for live cell analysis. The cells were treated with the appropriate concentration of recombinant proteins and YO-PRO (Thermo Fisher Scientific, diluted at 1:2000) and incubated at 37°C. Time-lapse confocal microscopy was performed using a NIKON A1R HD25 confocal microscope equipped with a 100× oil immersion objective. Images were captured at the indicated time points to observe the dynamic changes in the cells.

#### Western blotting

All antibodies used in the study are summarized in Table S2. Cell lysates were obtained using Minute^TM^ Total Protein Extraction Kit for Animal Cultured Cells/Tissues (Invent Biotech) according to the manufacturer’s instructions. After resuspension using 2× Laemmli loading buffer, protein samples were fractionated by SDS-PAGE and transferred to a PVDF membrane. The membranes were incubated overnight with primary antibodies, then with the corresponding peroxide-labeled IgG. Finally, enhanced chemiluminescence reagent was used to visualize the results.

#### Data analysis

Images were processed with NIS-Elements AR Analysis (Nikon, Inc.) and Adobe Illustrator CC (Adobe Systems, Inc.). The fluorescence intensity was analyzed using Image J (National Institutes of Health) and the corresponding graphs were generated by GraphPad Prism 8.

## CONCLUSIONS

The utilization of phase-separation modalities in the development of novel therapeutic proteins represents a paradigm shift in the field of signal transduction. The phase-separated Tumor Killers (psTK) designed in this study demonstrated remarkable interreceptor clustering and activation, leading to potent and selective induction of apoptosis in tumor cells. By leveraging liquid-liquid phase separation (LLPS) and its ability to organize cell surface receptors, we have achieved precise control over interreceptor communications, resulting in a highly efficient signaling pathway. The broad-spectrum capacity of psTK in targeting a wide range of tumor cells further highlights their potential as promising candidates for targeted therapies against challenging tumors. These findings have great significance, as they provide powerful tools for systematically manipulating multi-receptor complexes and modulating signaling outputs. Moreover, the innovative strategy of harnessing LLPS in the design of therapeutic proteins presented here opens new avenues for the design and development of programmable therapeutic agents with enhanced specificity and efficacy.

## ACKNOWLEDGMENTS

This study was supported by grants from National Natural Science Foundation of China 32150023 (P.L.). We are grateful to SLSTU-Nikon Biological Imaging Center (Center of Pharmaceutical Technology, Tsinghua University, Beijing, China) for imaging support, and Ting Wang (High Throughput Screening Core Facility, Center of Pharmaceutical Technology, Tsinghua University, Beijing, China) for SPR assays. We also thank the Tsinghua University Branch of China National Center for Protein Sciences (Beijing) and Tsinghua University Technology Center for Protein Research for the Cell Function Analyzing Facility support.

## DATA AVAILABILITY

This study includes no data deposited in external repositories. Expanded View for this article is available online.

## CONFLICT OF INTEREST

All authors declare that they have no competing interests.

## Contributions

P.L.L. and W.F.X. conceived of the study. Y.Y.L. and Y.T.Z. performed the experiments. Y.Y.L. conducted the statistical analyses. Y.Y.L. wrote the paper. P.L.L. and W.F.X. revised the paper. All authors reviewed the manuscript.

## EXPANDED VIEW FIGURE LEGENDS

**Figure EV1.**
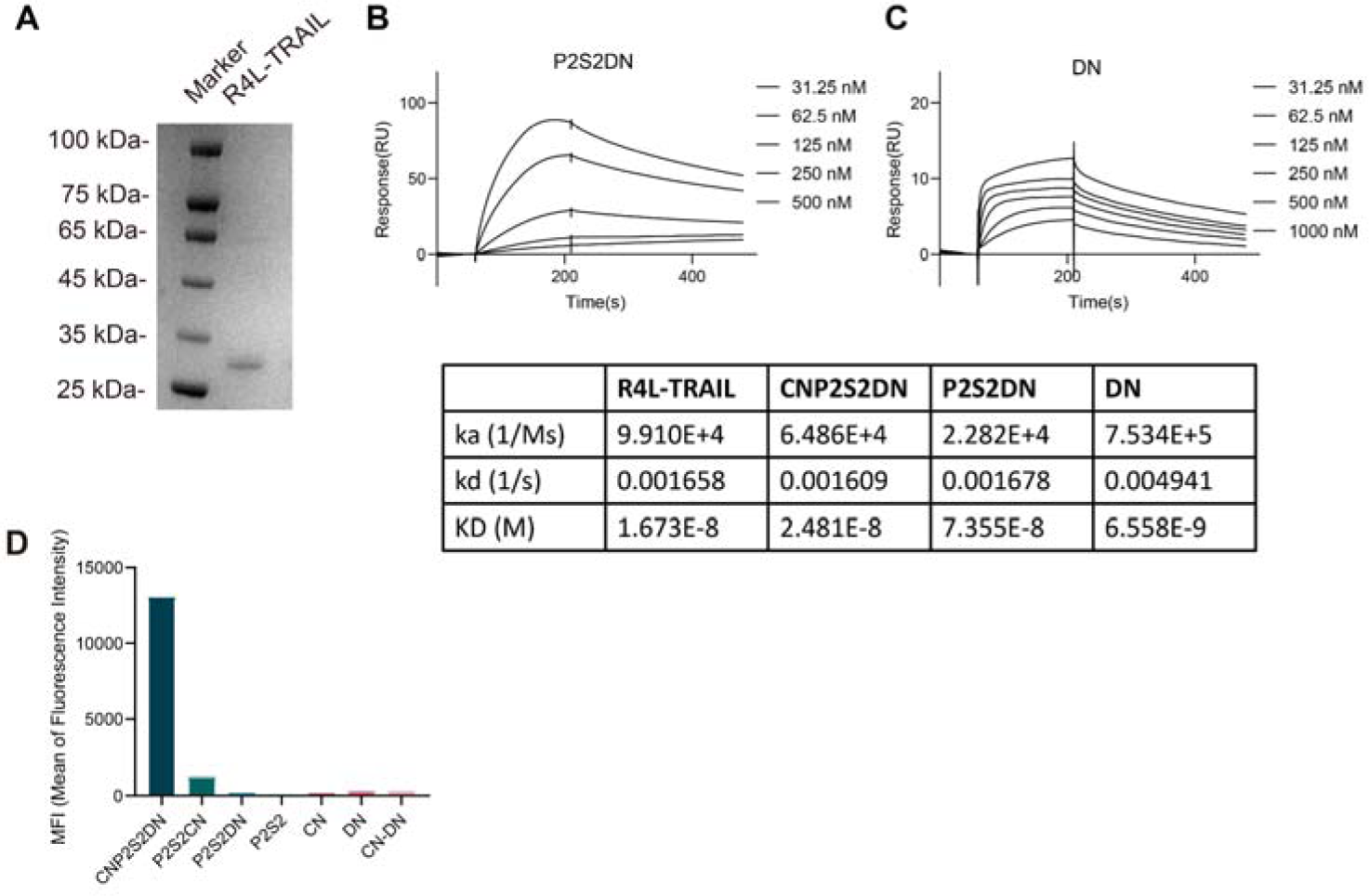
Analysis and Characterization of R4L-TRAIL and CN-P2S2-DN. **A** Analysis of R4L-TRAIL, fused with a C-terminal His_6_ tag, by SDS-PAGE. **B, C** Surface plasmon resonance (SPR) results showing the binding affinity of P2S2-DN and DN to DR5 extracellular domain (ECD). **D** Binding affinity of CN-P2S2-DN and control proteins to SJSA-1 cells assessed by flow cytometry.

**Figure EV2.**
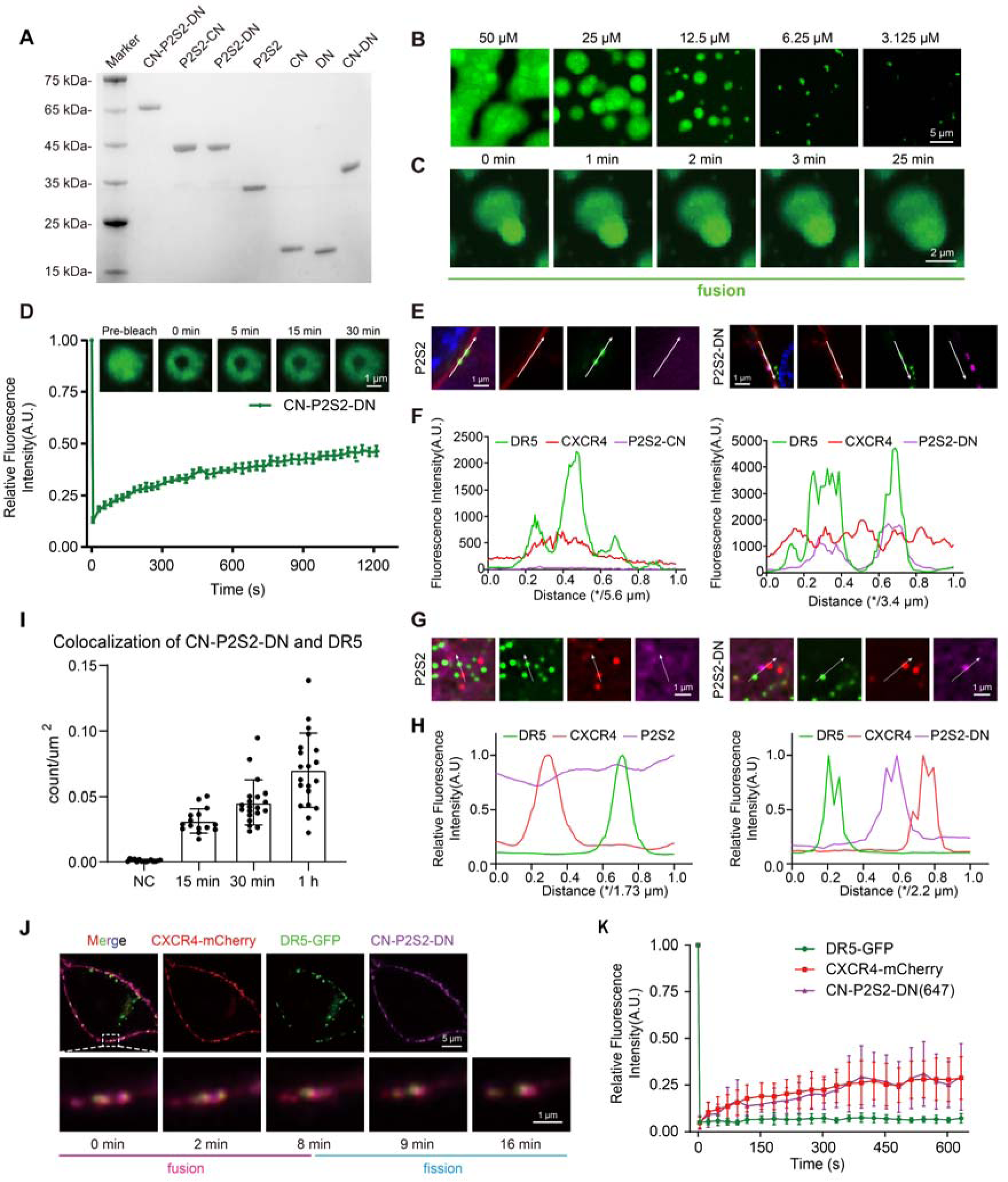
Characterization and Functional Analysis of CN-P2S2-DN: Phase Separation and Receptor Reorganization. **A** SDS-PAGE analysis of CN-P2S2-DN and corresponding truncations fused with a C-terminal His_6_ tag. **B** In vitro phase separation assay of CN-P2S2-DN labeled with Alexa 488. Confocal fluorescence imaging shows concentration-dependent formation of puncta. 10% PEG-8000 was added. Scale bar, 5 μm. **C** In vitro fusion process of puncta formed by 25 μM CN-P2S2-DN. **D** In vitro FRAP assay of puncta formed by 25 μM CN-P2S2-DN. The plot shows the time course of the recovery of CN-P2S2-DN. Data are representative of three independent experiments. Scale bar, 1 μm. **E** Confocal fluorescence imaging of HEK293T cells expressing DR5-EGFP and CXCR4- mCherry, treated with 200 nM P2S2-DN or P2S2 (Alexa 647). **F** Fluorescence intensity plots indicating that P2S2-DN and P2S2 cannot induce CXCR4 and DR5 colocalization. Fluorescence intensity profiles were analyzed along the white arrows in **(E)**. **G** Confocal imaging of immunofluorescence. SJSA-1 cells were treated with 100 nM P2S2 or P2S2-DN (Alexa 647), then fixed and incubated with respective primary antibodies and secondary antibodies. **H** Plots showing analysis of colocalization. Fluorescence intensities of P2S2/P2S2-DN, DR5, and CXCR4 were quantified along the white arrows in **(G)**. **I** Colocalization analysis of TIRF imaging statistics in SJSA-1 cells. The counts of colocalized CN-P2S2-DN and DR5 increased as the CN-P2S2-DN treatment time was extended. **J** HEK293T cells expressing DR5-EGFP and CXCR4-mCherry were treated with 200 nM P2S2-CN, P2S2-DN, or P2S2 (Alexa 647). Confocal fluorescence images are shown of the fusion and fission processes. **K** FRAP assay on the cell membrane. The plot shows the time course of the recovery of CN-P2S2-DN, DR5-GFP, and CXCR4-mCherry condensates on the SJSA-1 cell membrane. Time 0 indicates the time of the photobleaching pulse.

**Figure EV3.**
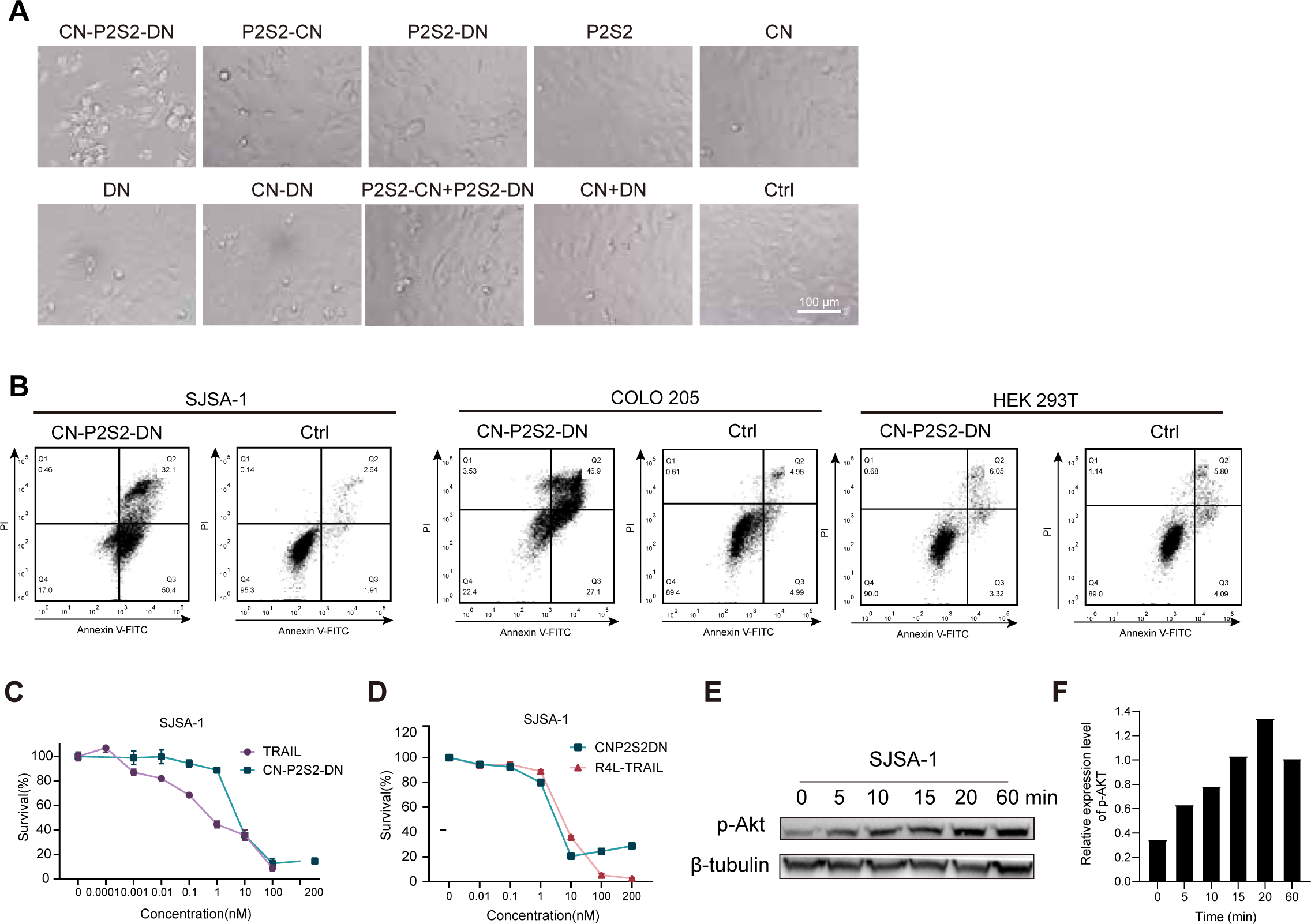
Additional Evidence of Selective Tumor-Specific Apoptosis Induction by CN-P2S2-DN. **A** Cellular imaging of SJSA-1 cells treated with 100 nM CN-P2S2-DN protein or control proteins for 3 hours, highlighting the selective induction of apoptosis in tumor cells. **B** Flow cytometry-based apoptosis analysis using the Annexin V-FITC/PI apoptosis analysis kit. SJSA-1, COLO 205, and HEK293T cells were treated with CN-P2S2-DN for 5 hours, and the percentage of cells undergoing late apoptosis and early apoptosis was determined. **C** Cell survival assessed by the CTG assay after 24-hour exposure of SJSA-1 cells to CN-P2S2-DN and TRAIL, demonstrating the comparable cytotoxicity of CN-P2S2-DN towards tumor cells. **D** Survival of SJSA-1 cells after 24 hours of treatment with the indicated concentrations of CN-P2S2-DN or R4L-TRAIL. **E** Western blot analysis showing enhanced activation of p-Akt in SJSA-1 cells within 60 minutes of treatment with 100 nM CN-P2S2-DN. This indicates the potential involvement of the Akt signaling pathway in the apoptotic response. **F** Quantitative analysis of p-AKT/β-tubulin band intensity in **(E)**.

**Figure EV4.**
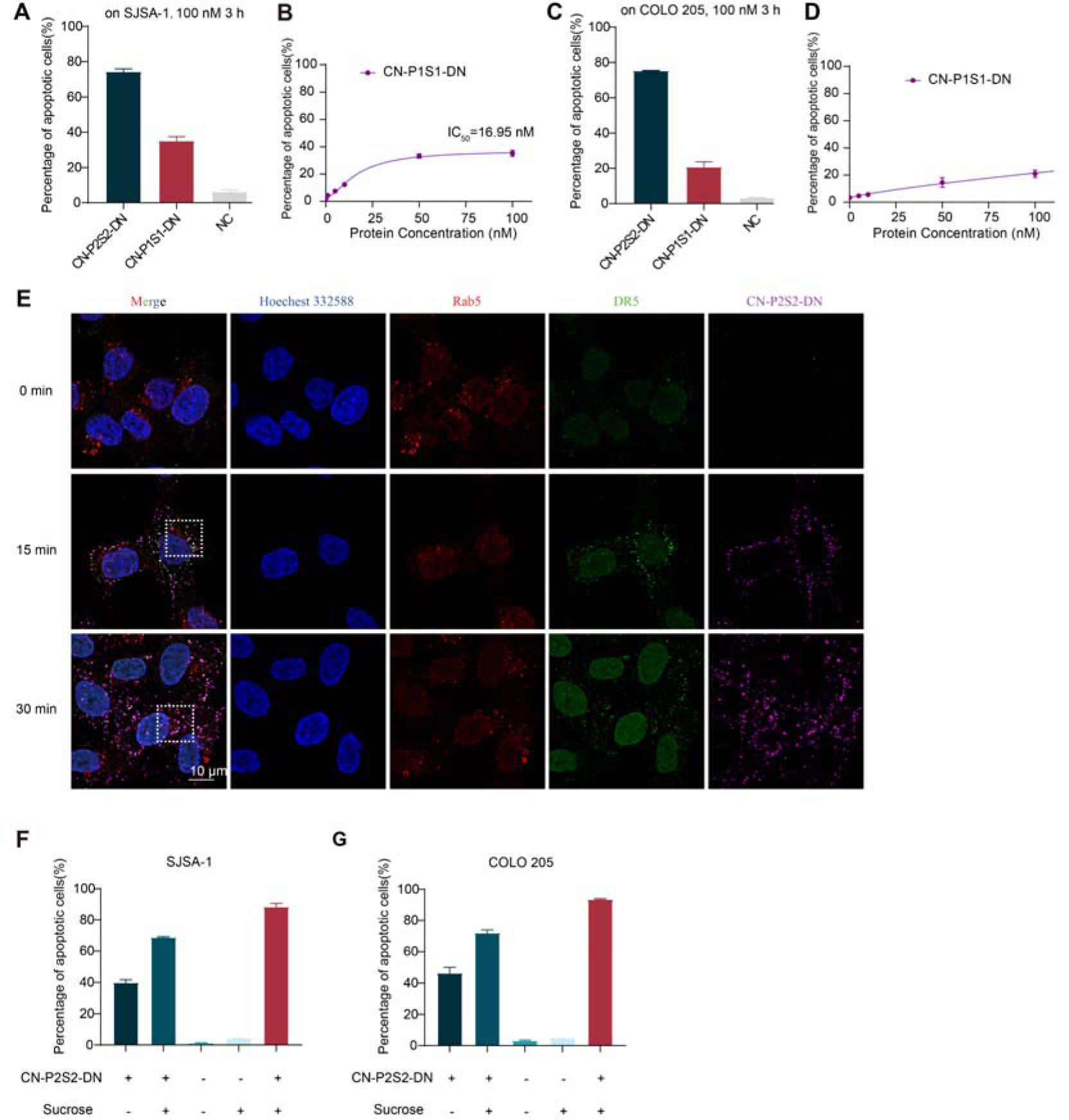
Subcellular Localization, Apoptosis Induction, and Valency-Dependent Analysis of CN-P2S2-DN. **A** Apoptosis induction analysis of low-valency CN-P1S1-DN and CN-P2S2-DN in SJSA-1 cells, demonstrating their respective abilities at varying doses. **B** Dose-response curve for CN-P1S1-DN treatment of SJSA-1 cells, emphasizing the valency-dependent apoptotic response. **C** Apoptosis induction analysis of low-valency CN-P1S1-DN and CN-P2S2-DN in COLO 205 cells, showing their effectiveness at different doses. **D** Dose-response curve for CN-P1S1-DN treatment of COLO 205 cells, highlighting the valency-dependent apoptotic response. **E** Dynamic subcellular localization analysis of CN-P2S2-DN in SJSA-1 cells, highlighting the entry of the ligands into the cells. Dotted boxes show the zoom region in Figure 4H. **F, G** Impact of sucrose on apoptosis induction by CN-P2S2-DN (10 nM) in SJSA-1 cells and COLO 205 cells.

**Figure EV5.**
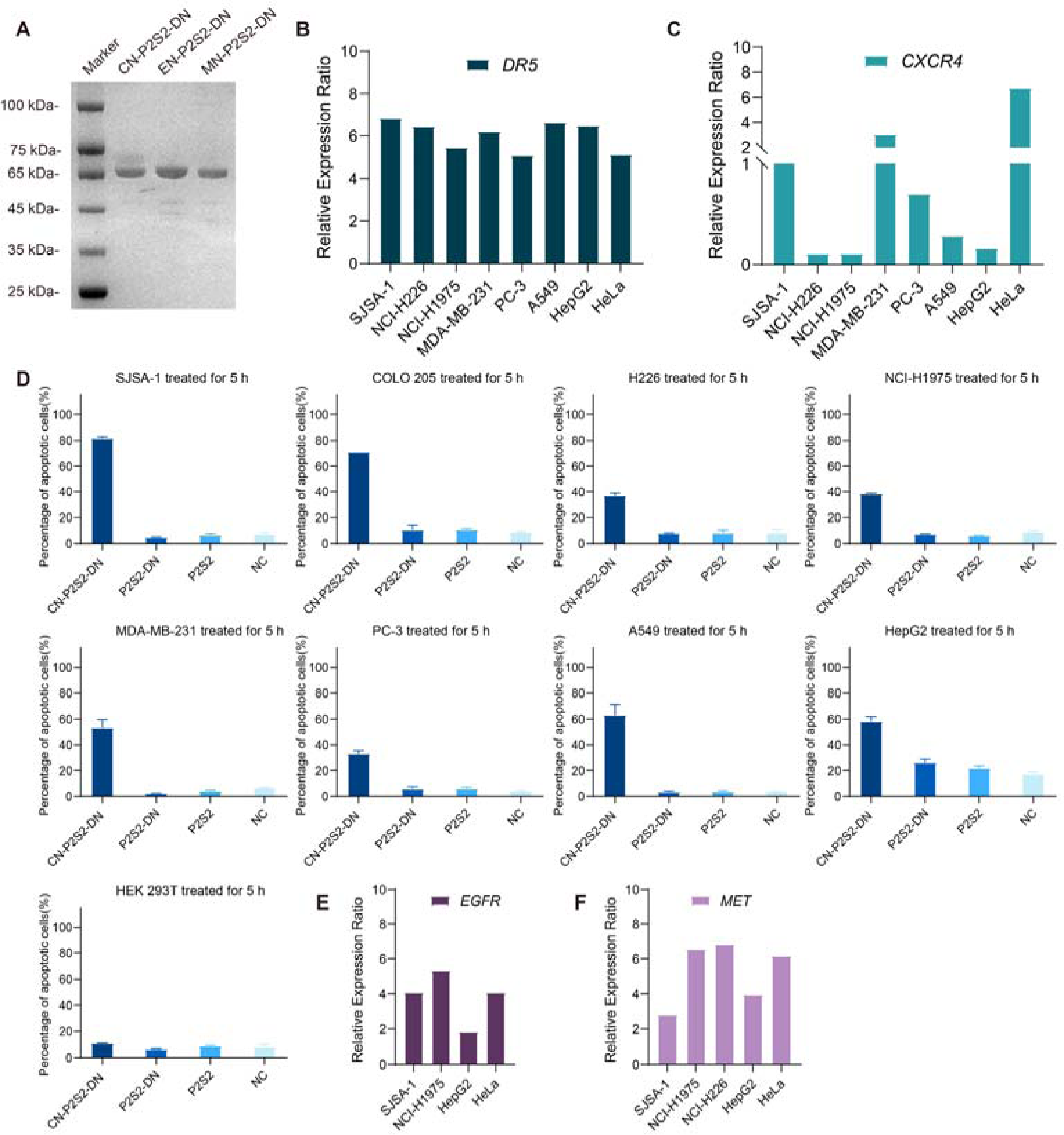
Characterization of Different psTK Constructs and Evaluation of Additional Target Cell Lines. **A** SDS-PAGE analysis of CN-P2S2-DN, EN-P2S2-DN, and MN-P2S2-DN fused with a C-terminal His_6_ tag. **B** Gene expression levels of DR5 in various cell lines, based on data from the DepMap database. **C** Gene expression levels of CXCR4 in various cell lines, based on data from the DepMap database. **D** Apoptosis analysis of SJSA-1, COLO 205, NCI-H226, NCI-H1975, MDAMB-321, PC-3, A549, HepG2, and 293T cells treated with CN-P2S2-DN or control proteins (200 nM) for 5 hours. **E** Gene expression levels of EGFR and MET in various cell lines, based on data from the DepMap database. **F** Gene expression levels of EGFR and MET in various cell lines, based on data from the DepMap database.

